# Integrative Functional Network Interactions Underlie the Association between Physical Activity and Cognition in Neurodegenerative Diseases

**DOI:** 10.1101/542936

**Authors:** Chia-Hao Shih, Miriam Sklerov, Nina Browner, Eran Dayan

## Abstract

Physical activity (PA) has preventive and possibly restorative effects in aging-related cognitive decline, which relate to intrinsic functional interactions (functional connectivity, FC) in large-scale brain networks. Preventive and ameliorative effects of PA on cognitive decline have also been documented in neurodegenerative diseases, such as Parkinson’s disease (PD). However, the neural substrates that mediate the association between PA and cognitive performance under such neurological conditions remain unknown. Here we set out to examine if the association between PA and cognitive performance in PD is mediated by FC in large-scale sensorimotor and association brain networks. Data from 51 PD patients were analyzed. Connectome-level analysis based on a whole-brain parcellation showed that self-reported levels of PA were associated with increased FC between, but not within the default mode (DMN) and salience networks (SAL) (p < .05, false discovery rate corrected). Additionally, multiple parallel mediation analysis further demonstrated that FC between left lateral parietal nodes in the DMN and rostral prefrontal nodes in the SAL mediated the association between PA and executive function performance. These findings are in line with previous studies linking FC in large-scale association networks with the effects of PA on cognition in healthy aging. Our results extend these previous results by demonstrating that the association between PA and cognitive performance in neurodegenerative diseases such as PD is mediated by integrative functional interactions in large-scale association networks.

## Introduction

Physical activity has preventive and potentially restorative effects targeting cognitive decline in normal aging (1–3) and in neurodegenerative diseases (4–6). In older adults, observational studies have shown that higher levels of physical activity are associated with better cognitive performance (7). Physically active older adults show lower rates of cognitive decline (8). In fact, physical activity is associated with a lower risk of developing neurodegenerative diseases (9–12), and may even delay the progression of neurodegeneration (13). In Parkinson’s disease (PD), for example, physical activity has been shown to positively impact both motor and non-motor symptoms, including cognitive function, as well as overall quality of life (14–16). Still, the mechanisms by which physical activity may mitigate cognitive decline in neurodegenerative diseases such as PD remain incompletely understood (13, 17).

In older adults, the benefits of physical activity are not only manifested in terms of performance in neuropsychological tests (18–21), but are also expressed in measurable markers of brain health, assessed using brain imaging techniques. Examples of this are the findings of increased global gray matter volume (22, 23), hippocampal size (24), and alterations in white matter integrity (25) reported in older adults following exercise interventions. More recently, studies utilizing resting-state functional magnetic resonance imaging (rs-fMRI) in aging individuals have shown that physical activity alters intrinsic functional interactions (i.e., functional connectivity) in large-scale association networks such as the default mode network (DMN), the salience network (SAL), the dorsal attention network (DAN), and the frontoparietal network (FPN), which are known to be altered in elderly people (26, 27). Cardiovascular fitness, a marker of moderate-intensity physical activity, positively relates to functional connectivity in the DMN, SAL, DAN, and executive control (ECN) networks (28, 29). This association is linked, in turn, to cognitive performance (29). Furthermore, exercise interventions were shown to enhance functional connectivity in association networks (e.g, DMN and FPN) but not in sensorimotor networks (30).

While converging evidence suggest that the association between physical activity and cognition in normal aging is mediated by interactions within and between large-scale brain networks, it remains incompletely understood if similar links exist in neurodegenerative diseases such as PD. Similar to the aging literature, studies in PD have reported that functional connectivity, particularly in association networks, is negatively associated with cognitive performance (31–33). Moreover, longitudinal cognitive decline in PD is associated with diffuse alterations in functional brain networks (34). Although potential neuroprotective effects of physical activity have been widely documented in relation to neurodegenerative diseases such as PD (4, 5, 14, 17, 35–38), the neural substrates that mediate the association between physical activity and cognitive performance in PD remain unknown.

In the current study, we analyzed behavioral and neuroimaging data collected as part of the multi-site Parkinson’s Progression Markers Initiative (PPMI). Analyzing imaging, clinical and behavioral measures collected within the same session, we set out to examine if the association between physical activity and cognitive performance in patients with de novo PD is mediated by functional connectivity in large-scale brain networks (Figure 1). We anticipated that physical activity would be associated with functional connectivity within and between large-scale functional networks, particularly in association networks such as the DMN, SAL, DAN, and FPN, which are known to change with age and respond to exercise interventions. We also hypothesized that functional connectivity within and between these networks would mediate the relationship between physical activity and cognition, with our analysis focusing on cognitive domains including executive function (i.e., Symbol Digit Modalities Test) and verbal memory (i.e., Hopkins Verbal Learning Test), which are the most frequent impaired cognitive domains in de novo PD (39).

**Figure 1.**
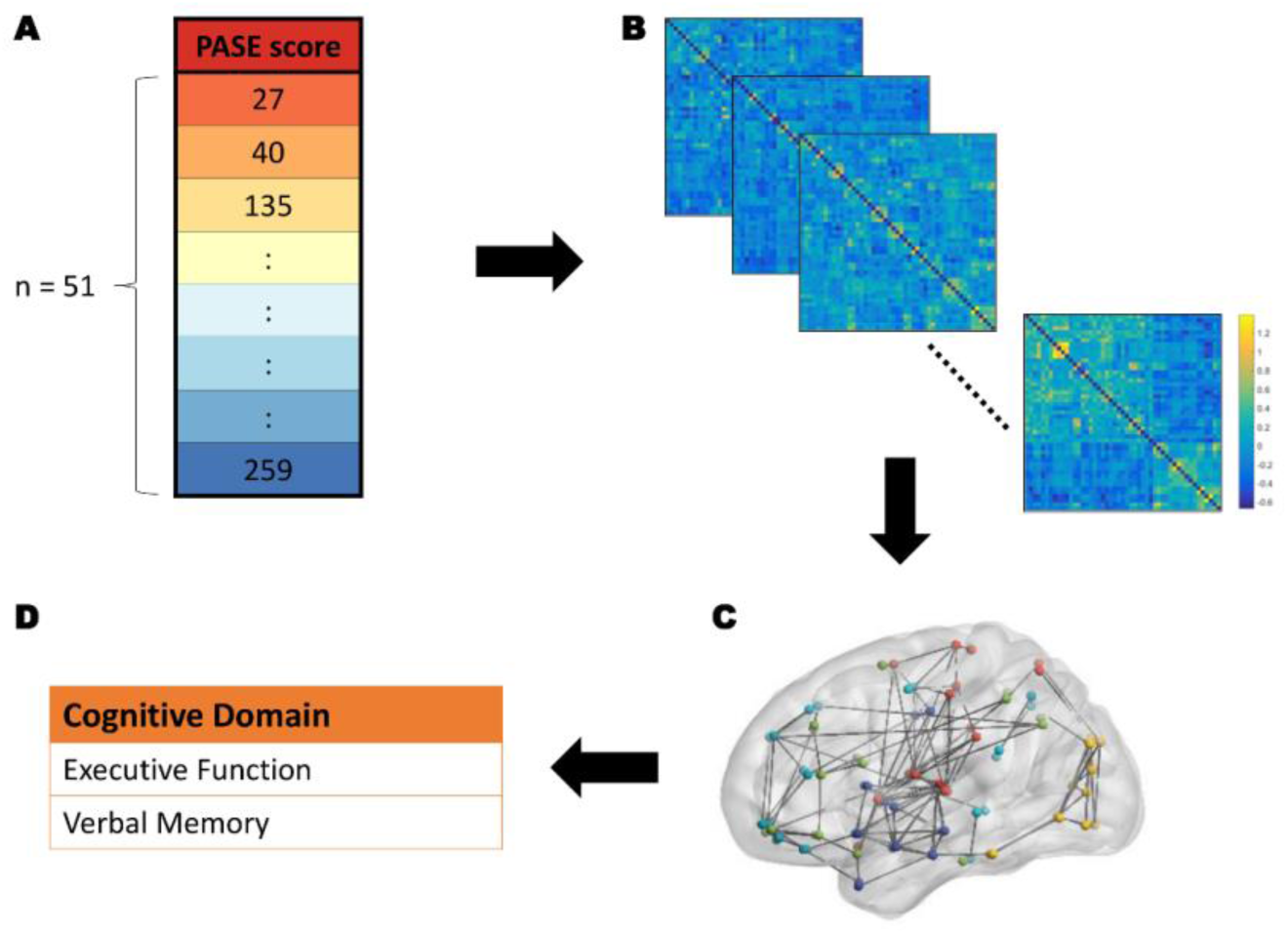
Analytical approach. First, we examine the association between physical activity, as assessed with the PASE questionnaire (A) and functional connectivity in large-scale brain networks (B and C). We then test whether the association between physical activity and cognitive performance (D) is mediated by functional connectivity in large-scale functional networks using a multiple parallel mediation model.

## Results

### Associations between Physical Activity and Functional Connectivity

We analyzed data from 51 patients with PD (mean age = 60.65 yrs, range: 40-79 yrs). Physical activity, measured with the validated Physical Activity Scale for Elderly (PASE) questionnaire (40, 41), was variable in our sample of subjects (Fig. 2a), with values varying by up to 62.93%. Across the entire sample of PD subjects, PASE scores were not correlated with subjects’ age, disease duration, disease severity as assessed with the Movement Disorder Society - Unified Parkinson’s Disease Rating Scale (MDS-UPDRS) part III (motor examination) in the “off” medications state, or with the patients’ Hoehn and Yahr disease stage (42) (Fig. 2b). No differences between these variables were found as a function of patients’ predominantly affected side (all *ps* > .05).

**Figure 2.**
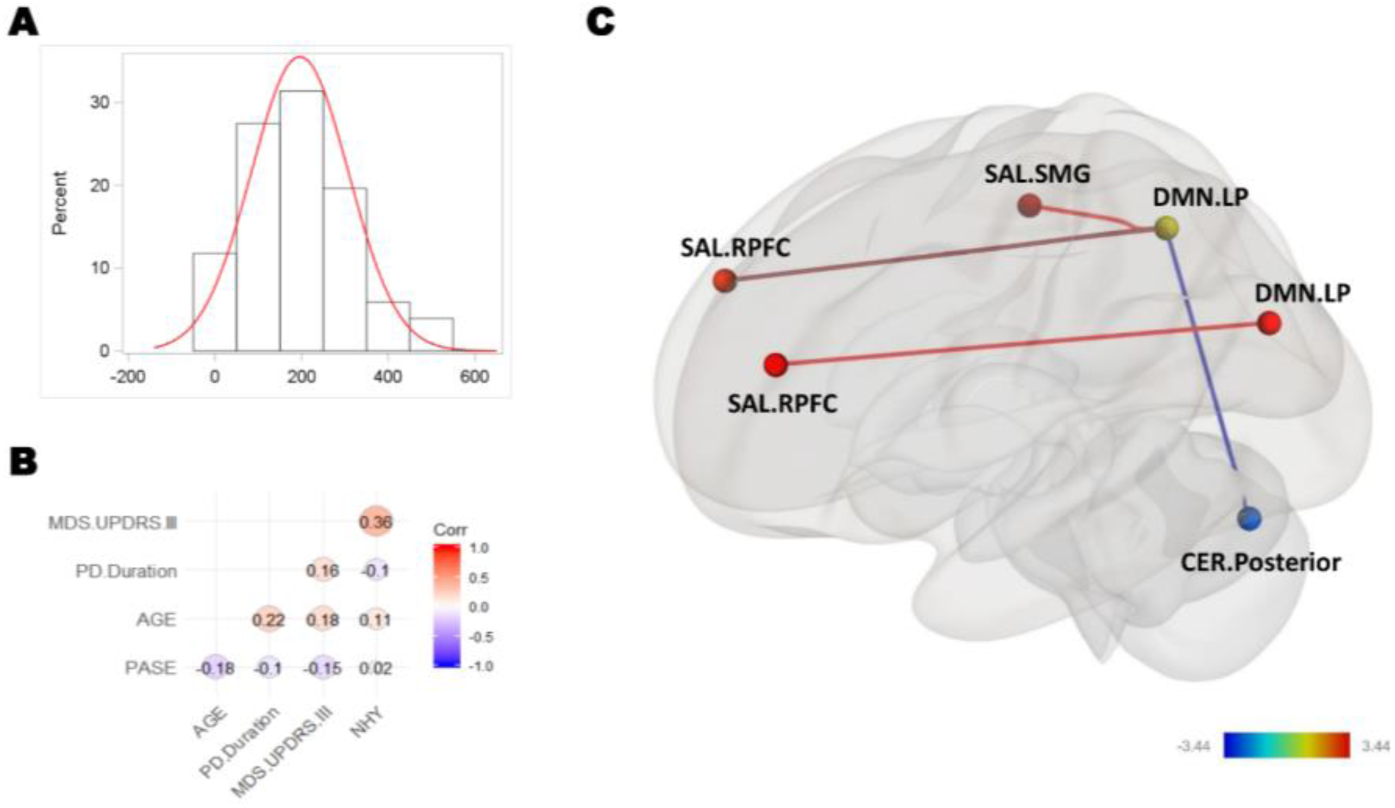
Association between physical activity and functional connectivity in large-scale functional networks. (A) Physical activity, assessed with the validated PASE questionnaire, was variable across the sample of PD patients. (B) PASE scores did not correlate significantly with other demographic and clinical characteristics. (C) Physical activity was significantly associated with functional connectivity between both left and right lateral parietal nodes in the default mode network (DMN.LP) and ipsilateral rostral prefrontal cortex nodes in the salience network (SAL.RPFC), as well as between right DMN.LP and right supramarginal gyrus nodes in the salience network (SAL.SMG). Physical activity was also negatively correlated with functional connectivity between right DMN.LP and posterior nodes in the cerebellar networks (CER.Posterior).

We first examined the association between physical activity scores and functional connectivity in large scale sensory, motor, and association networks (Supplemental Figure 1). Region of interest (ROI)-to-ROI analysis revealed associations between physical activity and functional connectivity, primarily in association networks (false-discovery rate [FDR] corrected for multiple comparisons) (Fig. 2c). Specifically, higher physical activity scores showed a significantly positive correlation with functional connectivity between left lateral parietal nodes in the DMN (DMN.LP) and ipsilateral rostral prefrontal cortex nodes in the SAL (SAL.RPFC), between right DMN.LP nodes and ipsilateral SAL.RPFC nodes, and between right DMN.LP and ipsilateral supramarginal gyrus nodes in the salience network (SAL.SMG). Thus, the results suggest that physical activity was mainly correlated with functional connectivity between, and not within large-scale association networks. In addition, physical activity was also negatively and significantly correlated with functional connectivity between right DMN.LP and the posterior nodes in the cerebellar networks (CER.Posterior).

The parcellation used in the analysis above is centered on large-scale sensory, motor, and association networks. Given that neurodegeneration in PD is primarily associated with progressive changes in basal ganglia circuitry (43) we next sought to examine whether functional connectivity in the basal ganglia correlated with physical activity scores. Based on a detailed atlas of the basal ganglia (44) (Supplemental Figure 2), this analysis revealed no significant associations between functional connectivity within the basal ganglia, or between the basal ganglia and the large-scale networks analyzed above, with physical activity in our sample of PD patients. We further evaluated whether the associations between physical activity and basal ganglia functional connectivity differed as a function of the patients’ predominantly affected side. Within the basal ganglia, functional connectivity contralateral to the patients’ predominantly affected side was not different compared to connectivity ipsilateral to their affected side (Fisher’s z test: z = 0.32, p = 0.75). Similar results were found for functional connectivity between basal ganglia and all other large-scale networks (Fisher’s z test: z = 0.01, p = 0.99).

### Mediating Role of Network Connectivity in the Association between Physical Activity and Cognitive Performance

We next tested the linkage between physical activity, functional connectivity in association networks, and cognitive performance. Specifically, we tested whether the association between physical activity and the patients’ performance in individual cognitive domains was mediated indirectly by the strength of functional connectivity in the edges connecting the DMN, SAL, and cerebellar networks identified above. Cognitive domains analyzed included executive function and verbal memory. Fitting the data with a multiple parallel mediation model (Figure 3A), the results indicate that the edges between left DMN.LP and ipsilateral SAL.RPFC significantly mediated the relationship between physical activity and executive function performance (β_DMN.LP-SAL.RPFC(left)_ = 0.0017, 95% CI [0.0004, 0.0036]) (Figure 3B and Supplemental Table 2). No other effects reached statistical significance for executive function or any of the other individual cognitive domains.

**Figure 3.**
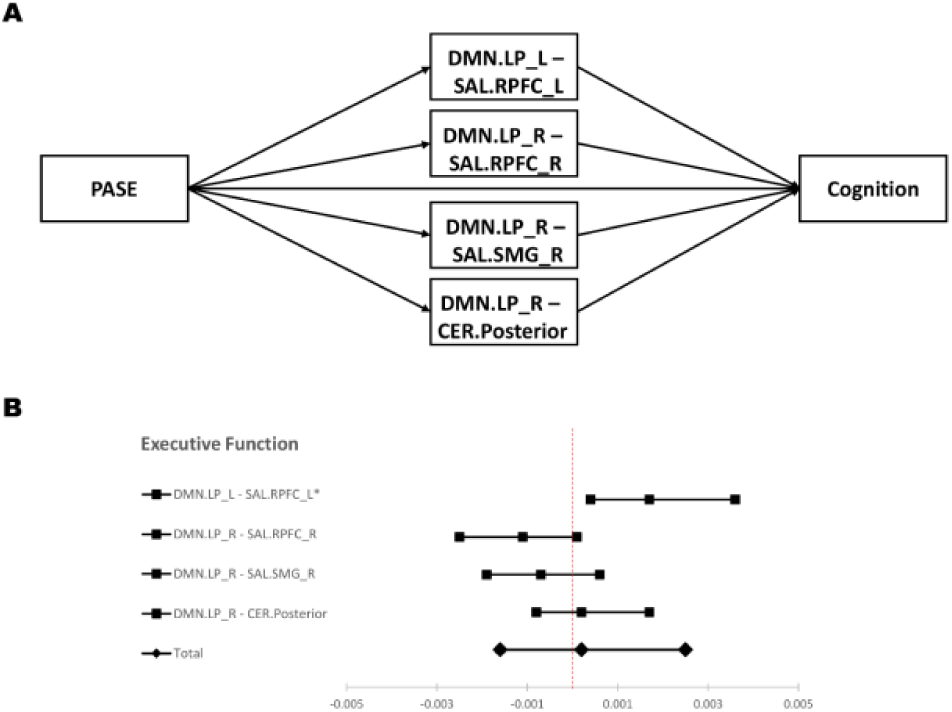
The linkage between physical activity, functional connectivity in large scale brain networks, and cognition. (A) Multiple parallel mediation model was used to examine the linkage between physical activity, functional connectivity, and subjects’ performance in individual cognitive domains. (B) 95% confidence interval (CI) of the indirect effects (individual and total) of functional connectivity on the association between physical activity and executive function. 95% CIs are displayed.

## Discussion

The beneficial effects of physical activity on cognitive function in individuals with neurodegenerative diseases such as PD have been well documented in the literature (35, 38, 45–48). However, the neural substrates that underlie the link between physical activity and cognitive performance in such neurological conditions received very little attention to date. Our results suggest that the association between physical activity and cognitive performance in de novo PD is mediated by integrative functional interactions in large-scale association networks, particularly involving interactions between the DMN and the SAL. Specifically, interactions between these two networks significantly mediated the relationship between physical activity and the patients’ executive function performance.

Results of the ROI-to-ROI analysis reported in the current study are in line with previous studies linking cardiovascular fitness and physical activity with functional interaction in large-scale brain networks, particularly association networks, including the DMN, SAL, DAN, and FPN, which are vulnerable to aging effects (28–30). In the current results, physical activity was also associated with an anti-correlation between lateral parietal nodes in the DMN and posterior nodes in the cerebellar network. While the interpretation of anti-correlations in the context of large-scale network interactions is under debate (49–52), anti-correlation between the DMN and other large-scale networks have been observed, both in the context of cognitive processing (53–55) and aging (56, 57). It has been suggested that the cerebellum is critical in age-related cognitive and motor performance (58) and plays crucial roles in pathological and compensatory effects in PD (59). Additional research is needed to better understand the role of interactions between these two networks in PD. No associations between physical activity and functional connectivity were found within the basal ganglia or between the basal ganglia and other large-scale brain networks. This may suggest that functional connectivity in association networks is more sensitive to variation in physical activity than functional connectivity in the basal ganglia. Indeed, previous studies have primarily associated physical activity with functional connectivity in the former, rather than the latter circuitry (28–30). Nevertheless, since our analyzed sample included patients with de novo PD, we cannot exclude that associations between physical activity and the basal ganglia may emerge later along the course of the disease.

Our findings demonstrate that functional connectivity between the DMN and the SAL mediates the association between physical activity and executive function in PD. Specifically, higher levels of physical activity were associated with increased functional connectivity between left lateral parietal nodes in DMN and ipsilateral rostral prefrontal cortex nodes in the SAL, which in turn, were associated with better executive function performance. These findings are largely consistent with previous findings in healthy older adults, where it was shown that cardiovascular fitness, a common marker of physical activity, was associated with increased functional connectivity in association networks including DMN, SAL, DAN, and ECN (28, 29), with functional connectivity in the DMN specifically showing an association with cognition (29). Other studies in older adults have also shown that exercise interventions lead to increased functional connectivity within and between association networks including the DMN and the SAL, which are associated with task performance in executive function (30, 60). Moreover, a linkage between fitness, network connectivity, and cognitive performance has also been documented in healthy older adults (29).

Our results support an association between physical activity and functional connectivity between but not within large-scale functional network, thus suggesting that integrative interactions underlie this link. Integrative functional network interactions have been suggested to provide an efficient way for information processing (61) and may be particularly critical for higher-order cognitive processes such as executive function (e.g., working memory) (62). Aberrant inter-network functional connectivity between the DMN and the SAL were reported in older adults with or without cognitive impairment (27, 63–65). Moreover, functional connectivity between these two networks has been linked to executive function performance (66). PD patients show impairment on a variety of cognitive domains (67, 68), with executive function being among the most frequent among these domains (69–71). Additionally, the literature suggests that executive function may benefit most strongly from physical activity (72–74). Together, these findings may explain the mediational link between physical activity, integrative functional connectivity between the DMN and the SAL, and executive function in PD reported in the current study.

Several limitations should be noted. First, the sample used in the current study was composed of de novo PD patients. Therefore, we cannot generalize our findings to other disease stages in PD. Second, due to the nature of the observational study design, we are not able to assess causal inference. Hence, future research with more appropriate designs (e.g., randomized controlled trials) is warranted to obtain more causal conclusions. Lastly, we did not analyze data from age-matched controls (see Methods) thus our results do not currently allow to make any inferences about differences between mechanisms operating in PD patients, relative to controls.

In summary, we report that physical activity in PD is associated with the strength of functional interactions between the DMN and the SAL. We in addition report that interactions between the DMN and the SAL mediate the relationship between physical activity and cognition in PD. The results suggest that integrative interactions among association networks, but not within or between the sensorimotor network and the basal ganglia, may underlie the link between physical activity and cognition in PD. Future research with robust study designs and diverse study samples will be needed to further understand the ameliorative role of physical activity in PD and other neurodegenerative diseases.

## Methods

### Participants

All data were obtained from the Parkinson’s Progression Markers Initiative (PPMI), a multi-center, observational study (75). Detailed information regarding PPMI’s mission, study design, inclusion and exclusion criterion, and data acquisition can be found online (www.ppmi-info.org). Research protocols were approved by the Institutional Review Boards of each participating centers, and written informed consent was provided by all participants before enrollment. Data from all PD patients who had concurrent (i.e., at the same visit) resting-state fMRI scans and physical activity measures were used in the current study (n=54, data downloaded on April 20, 2018). Data from 3 subjects were excluded from the sample due to excessive head motion during the MRI scan (i.e., > 50% of volumes detected as outliers), leaving 51 PD patients in the final analyzed sample. Participants’ demographic and clinical information is presented in Table 1. Since the PPMI did not have a sufficient number of healthy control subjects with neuroimaging and concurrent physical activity measures (n = 7), our analysis is based on data from the PD cohort alone.

**Table 1.** Demographic and clinical characteristics of the current sample.

| **<u>Demographic and clinical information</u>** |  |  |  |  |  |  |  |
| --- | --- | --- | --- | --- | --- | --- | --- |
|  | Age<br>(yr) | Sex<br>(F:M) | PASE | MDS-UPDRS III | H&Y Stages | PD duration<br>(mo) | Laterality<br>(L:R:S) |
| N | 51 | 13:38 | 51 | 49 | 49 | 51 | 20:29:2 |
| Mean | 60.65 | -- | 195.92 | 23.41 | 1.84 | 26.51 | -- |
| SD | 10.63 | -- | 112.31 | 12.66 | 0.37 | 14.38 | -- |
| **<u>Cognitive Outcomes</u>** |  |  |  |  |  |  |  |
|  | SDMT | HVLT<br>Total | HVLT<br>Delay | HVLT<br>Retention | HVLT<br>Recognition |  |  |
| N | 49 | 49 | 49 | 49 | 49 |  |  |
| Mean | 43.13 | 51.82 | 50.57 | 49.63 | 52.41 |  |  |
| SD | 9.69 | 12.72 | 11.73 | 10.27 | 11.03 |  |  |

*Note.* HVLT = Hopkins Verbal Learning Test; H&Y Stages = Hoehn and Yahr stages; Laterality = predominately affected side; MDS-UPDRS III = The Movement Disorder Society-Sponsored Revision of the Unified Parkinson's Disease Rating Scale part III; PASE = Physical Activity Scale for the Elderly; PD = Parkinson's disease; SDMT = Symbol Digit Modalities Test.

### Behavioral assessments

#### Physical activity

The Physical Activity Scale for the Elderly (PASE) (40) questionnaire was used to assess levels of physical activity. The PASE is a valid and reliable instrument, collected as part of the PPMI, which was specifically designed to assess both leisure and household physical activity in the older population. PASE scores show high correlations with objective measures of physical activity such as energy expenditure (76) and data derived from a portable accelerometers (77).

#### Cognitive function

Several neuropsychological tests are administered in PPMI. However, in the current study, we focus on cognitive domains of executive function and verbal memory based on previous research showing that these two are the most frequently impaired cognitive domains in PD (39). Executive function was assessed with the Symbol Digit Modalities Test (78) and verbal memory was tested with the Hopkins Verbal Learning Test – Revised (79, 80). For each individual participant, standardized scores were generated from the individual neuropsychological tests and composite scores in the domain of verbal memory were obtained by averaging scores from 4 subtests (81).

### MRI data acquisition

Imaging data were acquired with 3T Siemens scanners (Trio or Prisma). Structural 3D images were acquired using a magnetization-prepared rapid gradient echo sequence (echo time = 2.98 ms, repetition time = 2300 ms, flip angle = 9°, field of view = 256 x 256 mm, and voxel size = 1mm^3^). Resting-state fMRI scans were acquired using an echo-planar imaging sequence (echo time = 25 ms, repetition time = 2400 ms, flip angle = 80°, field of view = 222 x 222 mm, voxel size = 3.25 mm^3^, 40 axial slices in ascending order, slice thickness = 3.25 mm, and total number of scans = 210). During the resting-state fMRI scans, participants were instructed to rest quietly with their eyes open and attempt not to fall asleep. More details regarding imaging acquisition can be found online (www.ppmi-info.org).

### Data analysis

Imaging data analysis was performed in Matlab R2016b (MathWorks, Natick, MA) using the CONN toolbox (82), version 17. Brain images were first preprocessed including functional realignment and unwarping, slice-time correction, segmentation of gray matter, white matter, and cerebrospinal fluid (CSF) and normalization to the Montreal Neurological Institute template. Functional outlier detection was implemented with the ART-based scrubbing method, based on a 2 mm subject motion threshold and a global signal threshold of Z = 9. Nuisance variables, including 6 motion realignment parameters and their first-order derivatives, and signals from the segmented white matter and CSF were regressed out of the signal. Outlier volumes detected in the scrubbing procedure were left of the analysis as well. The data were subsequently band-pass filtered at 0.008 to 0.09 Hz.

A region of interest (ROI)-to-ROI analysis was then conducted to examine the effects of physical activity on functional connectivity in PD. 32 ROIs constituting the DMN, SAL, DAN, FPN, sensorimotor, visual, language, and cerebellar networks were first defined based on a network parcellation provided with the CONN toolbox. This parcellation is based on an independent component analysis of 497 resting-state fMRI scans from the Human Connectome Project. Six ROIs from the Harvard-Oxford atlas (i.e., left and right caudate, putamen, and thalamus) and 11 additional ROIs were defined based on age-specific atlas of the basal ganglia (44). Peak coordinates of all ROIs are list in Supplemental Table 1. To evaluate the association between physical activity and functional connectivity across all participants, second level analyses were implemented using an Analysis of Covariance (ANCOVA). ROI-to-ROI results are reported when significant at a level of p < .05, FDR-corrected. In all statistical tests, missing values (see Table 1) were excluded from the analysis.

Other statistical analyses were performed using SAS 9.4 (SAS Institute Inc., Cary, NC) and the PROCESS macro developed by Hayes (83). Multiple parallel mediation analysis was performed to examine whether functional connectivity in large-scale functional networks mediated the relationship between physical activity and each cognitive domains. In order to test the statistical significance of the indirect (i.e., mediation) effects, 95% confidence intervals (CI) were established by using a bootstrapping approach with 5,000 samples. Statistical significance in these tests is obtained whenever the 95% CI falls outside the value of 0.

## Acknowledgement

Data were obtained from the Parkinson’s Progression Markers Initiative (PPMI) database. For up-to-date information on the study, visit www.ppmi-info.org.

PPMI – a public-private partnership – is funded by the Michael J. Fox Foundation for Parkinson’s Research and funding partners, including Abbvie, Avid Radiopharmaceuticals, Biogen, Bristol-Myers Squibb, Covance, GE healthcare, Genentech, GlaxoSmithKline, Lilly, Lundbeck, Merck, Meso Scale Discovery, Pfizer, Piramal, Roche, Servier, and UCB.

## Supplementary Materials for

**Supplemental Figure 1.**
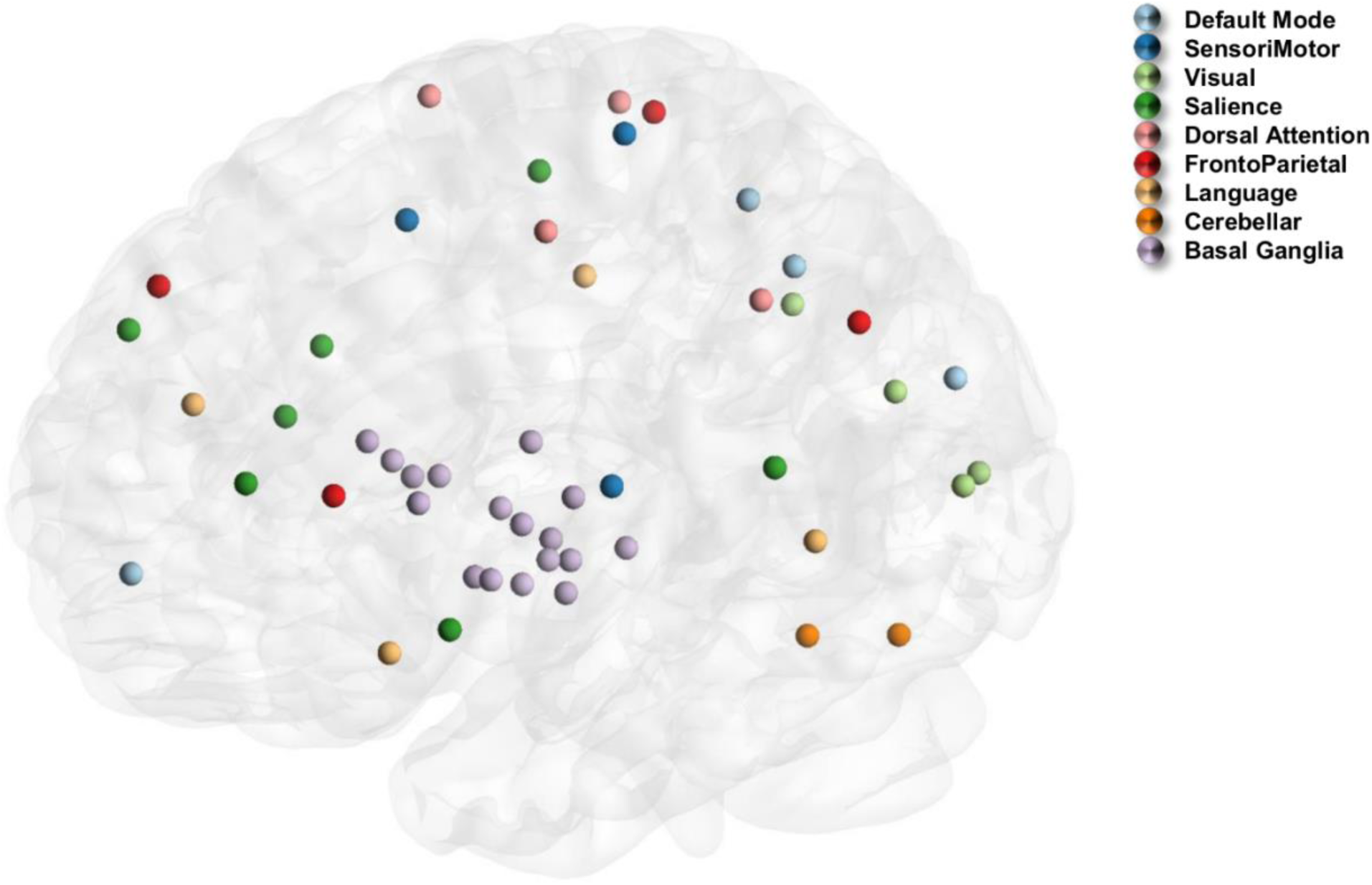
All nodes (color-coded based on networks) included in the functional connectivity analysis.

**Supplemental Figure 2.**
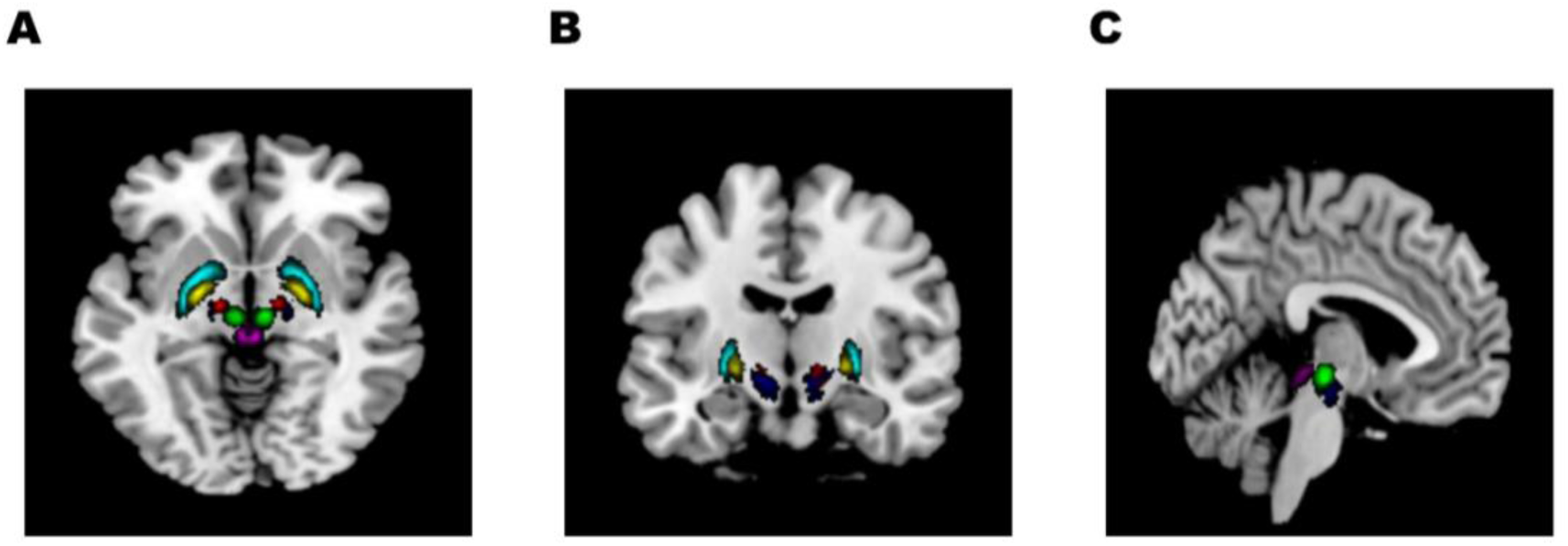
Detailed atlas of the basal ganglia from (A) Axial view, (B) Coronal view, and (C) Sagittal view.

**Supplemental Table 1.** Descriptions and coordinates of the region of interest (ROI) used in the current study.

| ROI | ROI label | ROI description | Laterality | NMI | Coordinate |  |
| --- | --- | --- | --- | --- | --- | --- |
|  |  |  |  | X | Y | Z |
| Functional Networks |  |  |  |  |  |  |
| Default Mode | MPFC | Medial prefrontal cortex | -- | 1 | 55 | -3 |
| Default Mode | LP | lateral parietal | L | -39 | 77 | 33 |
| Default Mode | LP | lateral parietal | R | 47 | -67 | 29 |
| Default Mode | PCC | posterior cingulate cortex | -- | 1 | -61 | 38 |
| SensoriMotor | Lateral | lateral portions | L | -55 | -12 | 29 |
| SensoriMotor | Lateral | lateral portions | R | 56 | -10 | 29 |
| SensoriMotor | Superior | superior portions | -- | 0 | -31 | 67 |
| Visual | Medial | medial portions | -- | 2 | -79 | 12 |
| Visual | Occipital | occipital portions | -- | 0 | -93 | -4 |
| Visual | Lateral | lateral portions | L | -37 | -79 | 10 |
| Salience | ACC | anterior cingulate cortex | -- | 0 | 22 | 35 |
| Salience | AInsula | anterior insula | L | -44 | 13 | 1 |
| Salience | AInsula | anterior insula | R | 47 | 14 | 0 |
| Salience | RPFC | rostral prefrontal cortex | L | -32 | 45 | 27 |
| Salience | RPFC | rostral prefrontal cortex | R | 32 | 46 | 27 |
| Salience | SMG | supramarginal gyrus | L | -60 | -39 | 31 |
| Salience | SMG | supramarginal gyrus | R | 62 | -35 | 32 |
| Dorsal Attention | FEF | frontal eye fields | L | -27 | -9 | 64 |
| Dorsal Attention | FEF | frontal eye fields | R | 30 | -6 | 64 |
| Dorsal Attention | IPS | intraparietal suclus | L | -39 | -43 | 52 |
| Dorsal Attention | IPS | intraparietal suclus | R | 39 | -42 | 54 |
| FrontoParietal | LPFC | lateral prefrontal cortex | L | -43 | 33 | 28 |
| FrontoParietal | PPC | posterior parietal cortex | L | -46 | -58 | 49 |
| FrontoParietal | LPFC | lateral prefrontal cortex | R | 41 | 38 | 30 |
| FrontoParietal | PPC | posterior parietal cortex | R | 52 | -52 | 45 |
| Language | IFG | inferior frontal gyrus | L | -51 | 26 | 2 |
| Language | IFG | inferior frontal gyrus | R | 54 | 28 | 1 |
| Language | pSTG | posterior superior temporal gyrus | L | -57 | -47 | 15 |
| Language | pSTG | posterior superior temporal gyrus | R | 59 | -42 | 13 |
| Cerebellar | Anterior | anterior portions | -- | 0 | -63 | -30 |
| Cerebellar | Posterior | posterior portions | -- | 0 | -79 | -32 |
| Basal Ganglia |  |  |  |  |  |  |
|  | Thalamus | Thalamus | L | 11 | -18 | 7 |
|  | Thalamus | Thalamus | R | -10 | -19 | 6 |
|  | Caudate | Caudate | L | 13 | 10 | 10 |
|  | Caudate | Caudate | R | -13 | 9 | 10 |
|  | Putamen | Putamen | L | 25 | 2 | 0 |
|  | Putamen | Putamen | R | -25 | 0 | 0 |
|  | GPe | external globus pallidus | L | -19 | 1 | -2 |
|  | GPe | external globus pallidus | R | 20 | 0 | -1 |
|  | GPi | internal globus pallidus | L | -20 | -7 | -4 |
|  | GPi | internal globus pallidus | R | 21 | -5 | -2 |
|  | PAG | periaqueductal gray | -- | -1 | -31 | -9 |
|  | RN | red nucleus | L | -5 | -20 | -8 |
|  | RN | red nucleus | R | 6 | -20 | -9 |
|  | SN | substantia nigra | L | -9 | -18 | -12 |
|  | SN | substantia nigra | R | 13 | -17 | -9 |
|  | STN | subthalamic nucleus | L | -12 | -14 | -4 |
|  | STN | subthalamic nucleus | R | 12 | -13 | -5 |

**Supplemental Table 2.** Coefficient estimation of the multiple parallel mediation model.

|  | Bias corrected bootstrapping CI |  |  |  |
| --- | --- | --- | --- | --- |
|  | Product of coefficients |  | 95% CI |  |
|  | Point estimate | SE | Lower | Upper |
| **Executive Function** |  |  |  |  |
| DMN.LP_L - SAL.RPFC_L* | 0.0017 | 0.0008 | 0.0004 | 0.0036 |
| DMN.LP_R - SAL.RPFC_R | -0.0011 | 0.0007 | -0.0025 | 0.0001 |
| DMN.LP_R - SAL.SMG_R | -0.0007 | 0.0006 | -0.0019 | 0.0006 |
| DMN.LP_R - CER.Posterior | 0.0002 | 0.0006 | -0.0008 | 0.0017 |
| Total | 0.0002 | 0.0010 | -0.0016 | 0.0025 |

*Note*<sup>1</sup>. \* indicates significant indirect effect.
*Note*<sup>2</sup>. CI = Confidence Interval; SE = Standard Error.

